# Probing direct interactions between nuclear proteins in cells with nxReLo

**DOI:** 10.1101/2025.07.14.664692

**Authors:** Sherif Ismail, Jana Kubíková, Maria Maichel, Peter Andersen, Mandy Jeske

## Abstract

Nearly all biological processes depend on protein-protein interactions (PPIs). While various methods exist to study these interactions, investigating those that involve nuclear proteins, including structurally complex proteins containing long disordered regions, remains a significant challenge. Here, we developed nxReLo, a simple and fast cell culture-based colocalization assay, designed to identify and characterize interactions between nuclear proteins. PIWI-interacting RNAs (piRNAs) safeguard germline genome integrity, and the PPIs that facilitate piRNA production are therefore essential for animal reproduction, yet remain incompletely understood. We used nxReLo to investigate interactions between members of the *Drosophila melanogaster* Rhino-Deadlock-Cutoff (RDC) complex and two associated components, Bootlegger and Moonshiner, a nuclear protein network required for piRNA expression. We demonstrate the utility of the nxReLo assay by systematically screening pairwise interactions within the RDC network and by assembling a multiprotein complex from multiple components. By combining nxReLo assays with AlphaFold structural prediction, we characterized the Cutoff-Deadlock and the Bootlegger-Deadlock complexes in detail, providing molecular and structural insights. Specifically, we refined the domains involved in the interaction and identified interface point mutations that interfered with complex formation, validating the predicted structures. In conclusion, nxReLo facilitates rapid and simple testing of direct interactions between nuclear proteins in a cellular context, which is particularly important when working with structurally challenging proteins or when established interaction assays prove unsuccessful.

## INTRODUCTION

All forms of life depend on the function of proteins. Central to these functions are interactions with small molecules, lipids, nucleic acids, and other proteins. The nucleus is a core cellular compartment in eukaryotes, encompassing complicated protein-protein interaction networks that serve diverse critical functions, including DNA replication and transcription, chromatin remodeling, epigenetic regulation, RNA processing, and ribosome biogenesis. To understand biological processes, the detection and characterization of protein-protein interactions (PPIs) is fundamental. Many techniques have been developed to study PPIs, each with its own advantages and limitations.

PPIs are commonly identified using mass spectrometry-based screening techniques, such as co-immunoprecipitation (co-IP), tandem affinity purification, and proximity labeling methods including BioID and APEX (1–7). These approaches typically yield a list of potential interaction partners, ranked by their relative abundance in the eluates. However, distinguishing direct binders from indirect associations often requires additional validation, frequently involving in vitro assays like GST pull-down assays, which depend on the availability of purified proteins. Obtaining soluble proteins from recombinant expression can be difficult, especially if the proteins contain a high degree of unstructured regions, as often observed for nuclear proteins (8, 9).

As an alternative to *in vitro* methods, interactions between nuclear proteins can be tested using cell-based assays (10–15), of which the most established is yeast-two-hybrid (Y2H) (16, 17). In Y2H, the PPI is detected by the activation of reporter genes. However, some proteins, such as transcription factors, can cause an interaction readout, even in the absence of a binding partner. Another limitation of the assay is that information on protein expression is not directly available but requires additional western blot analysis, which complicates the process when investigating many interactions. If the candidate proteins are poorly expressed or rapidly degraded in the cell, and the expression is not carefully evaluated, this can lead to the false assumption that an interaction is absent (false negative result).

We recently developed ReLo, a robust and rapid method to detect and characterize PPIs in a cellular context, particularly suited for structurally complex proteins that are difficult to obtain through recombinant expression (18). In the ReLo assay, a fluorescently tagged bait protein is anchored to the cytoplasmic side of the plasma membrane, while the prey protein, carrying a different fluorescent tag, remains unanchored. If the proteins interact, the prey relocalizes to the membrane alongside the bait. Relocalization is typically observed in over 90% of co-transfected, morphologically normal cells. Importantly, we have shown that interactions detected by ReLo are very likely direct and not mediated by endogenous cellular factors. While the assay has been successfully applied to interactions involving at least one cytoplasmic protein (19–22), testing two nuclear proteins may yield inconclusive results, as a nuclear prey may be absent from the cytoplasm and therefore fail to relocalize to the membrane (**Figure 1**).

**Figure 1.**
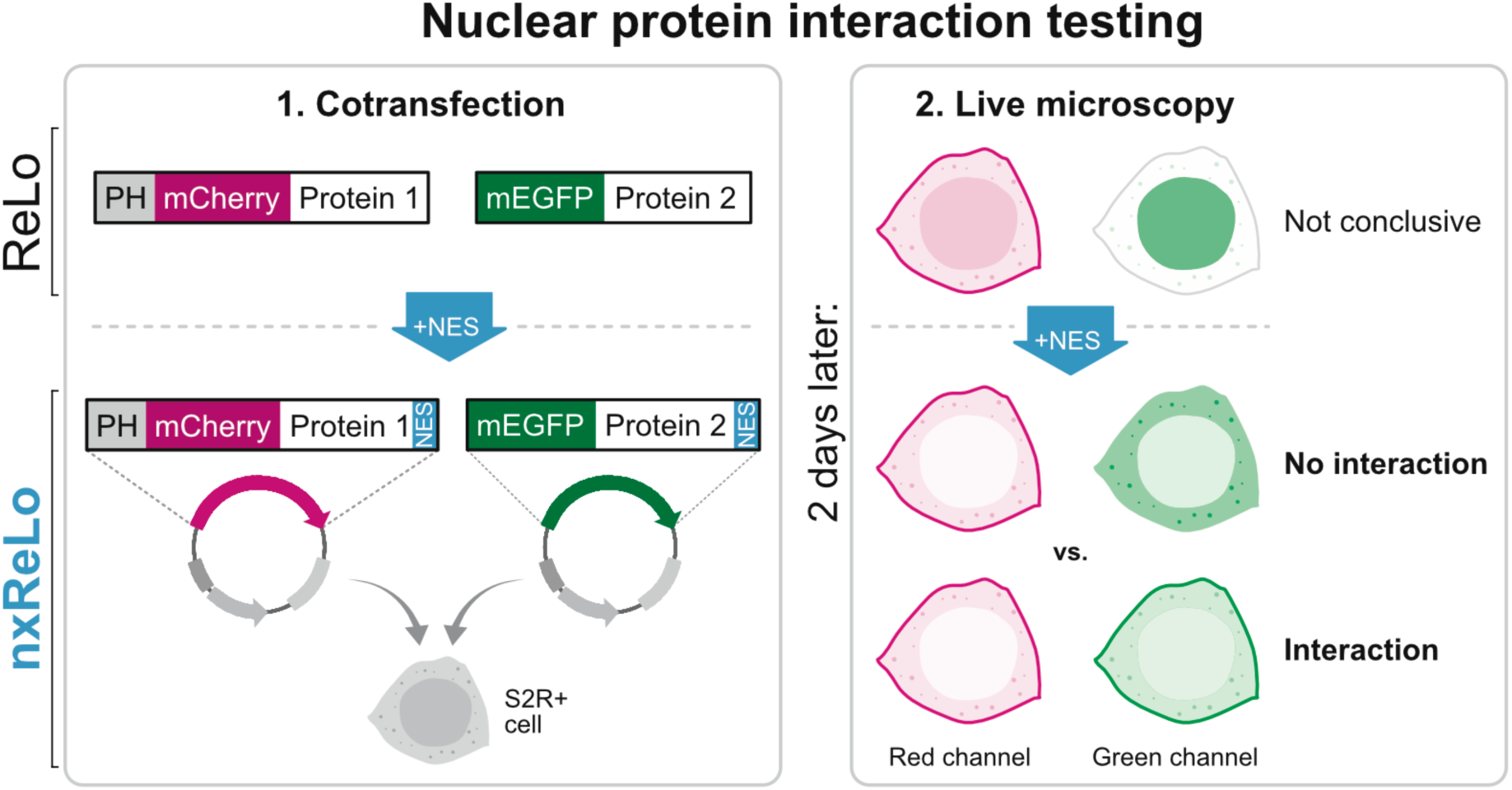
The nxReLo assay. Plasmids encoding fluorescently tagged protein 1 (bait) and protein 2 (prey), both fused to an NES, are cotransfected into S2R+ cells. Protein 1 is also fused to a plasma membrane anchoring domain (PH). The subcellular localization of the proteins is visualized by confocal microscopy after 48 hours post-transfection. In case of an interaction with protein 1, protein 2 is relocalized to the plasma membrane.

In this work, we developed <u>n</u>uclear e<u>x</u>port ReLo (nxReLo), a cell-based relocalization assay that allows the unambiguous study of interactions between two nuclear proteins (**Figure 1**). The nxReLo assay is an adaptation of the ReLo assay, in which the protein interaction partners are fused to a nuclear export signal (NES), leading to their export to the cytoplasm, and thus their presence in the same cellular compartment. As a proof of principle, we used nxReLo to investigate the interactions between the members of the nuclear *Drosophila melanogaster* Rhino-Deadlock-Cutoff (RDC) complex and two associated proteins, Bootlegger and Moonshiner. The RDC complex drives the non-canonical transcription of repeat-rich genomic loci to express Piwi-interacting RNAs (piRNAs) (23–28). piRNAs safeguard germline genome integrity by directing Argonaute-mediated silencing of transposons and other repetitive elements (recently reviewed in (29, 30). The PPIs facilitating piRNA production are thus essential for animal reproduction, but remain incompletely understood. By combining nxReLo with AlphaFold structure prediction, we investigated the interactions between Deadlock and Cutoff, as well as Deadlock and Bootlegger, two structurally uncharacterized complexes, and verified the predicted binding interfaces with mutational analysis. We have also utilized the potential of nxReLo for bridging interactions to describe the binding topology of the RDC network and to assemble a four-component complex. As such, we introduce nxReLo as an effective method to rapidly detect and characterize direct interactions between nuclear proteins.

## RESULTS

### Systematic pairwise nxReLo interaction screen of a nuclear five-component protein network

In the nxReLo assay, the bait protein is anchored to the plasma membrane via a pleckstrin homology (PH) domain and fused to a red fluorescent protein (mCherry). The prey protein remains unanchored and is fused to a green fluorescent protein (mEGFP). Both constructs also contain a modified NS2 nuclear export signal (NES; TSDEMTKKFGTLTI) (31) at their C-termini to promote export to the cytoplasm **(Figure 1)**. This sequence is referred to as NES throughout this study. The constructs are coexpressed in cultured *Drosophila* S2R+ cells, and protein localization is assessed by confocal fluorescence microscopy.

The RDC complex and its associated proteins Bootlegger and Moonshiner are expressed in *Drosophila* germline cells and are essential for piRNA transcription and export (23–27, 32–34). For simplicity, we refer to all five components as the RDC network in this study (**Figure 2A**). Previous studies, including X-ray crystallography, showed that Rhino, a heterochromatin protein 1 (HP1)/chromobox (CBX) variant, recognizes and binds to trimethylated lysine residues (K9) of histone 3 (H3) protein tails (H3K9me3) present at heterochromatic piRNA clusters (24, 28, 35), and also interacts with the adaptor protein Deadlock (24, 36). Y2H assays demonstrated that Deadlock interacts with Cutoff and Bootlegger (24, 33). Co-immunoprecipitation (co-IP) experiments further revealed that Deadlock co-purifies with Moonshiner, a protein involved in transcription initiation at piRNA clusters (23). However, while *D. melanogaster* Deadlock and Moonshiner did not interact in Y2H assays, their *D. simulans* orthologs did (22). An interaction between Cutoff and Rhino was suggested by co-IP (28, 32), but not detected in Y2H assays (22, 24). Because all components of the RDC network are localized in the nucleus *in vivo* (23–25, 32–34), and interaction data mostly from Y2H and co-IP studies are already available, this network represents a suitable test case for validating and applying the nxReLo assay.

**Figure 2.**
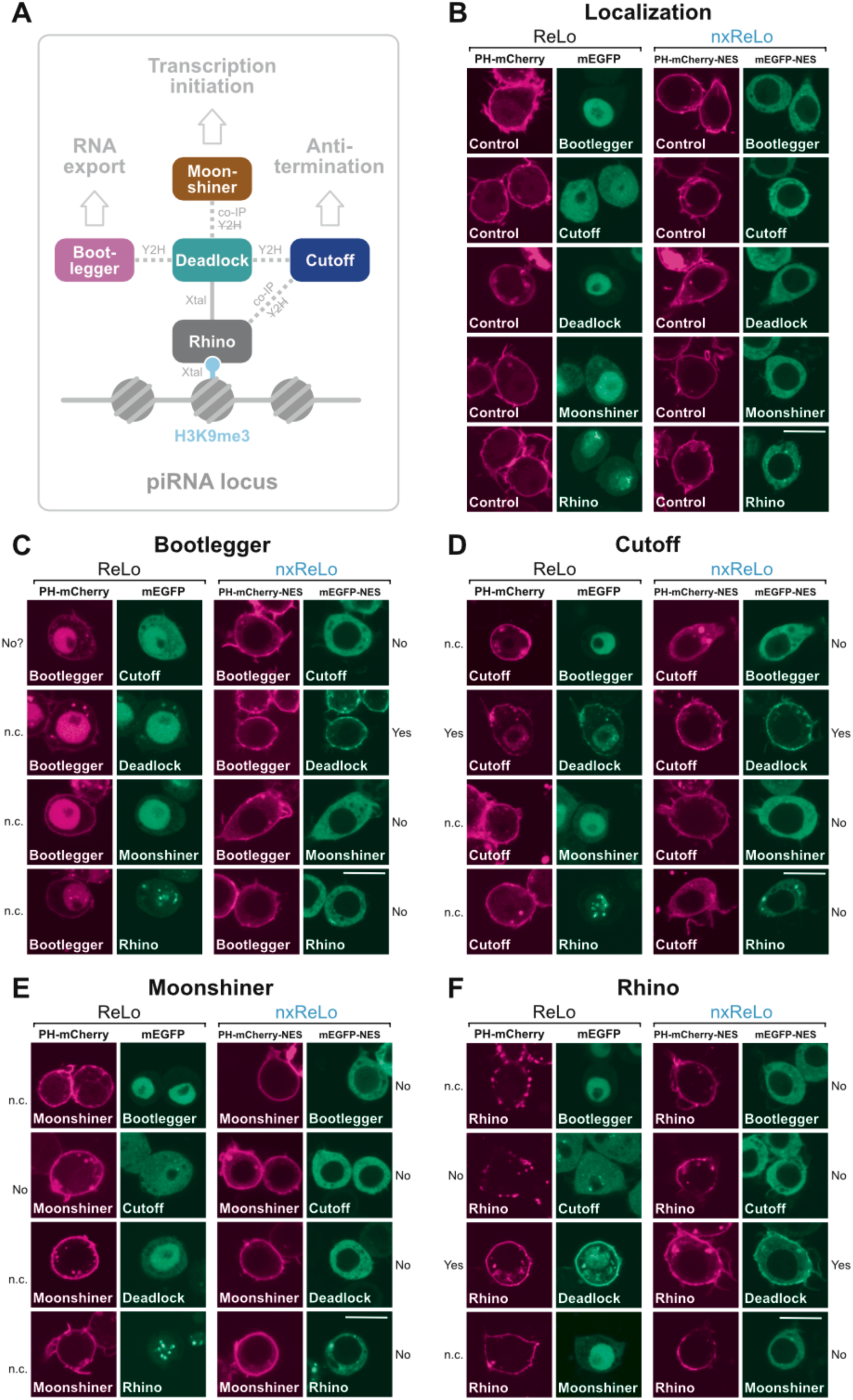
Pairwise interaction screen between members of the RDC network. (A) Schematic representation of the RDC network, its functions, and previously reported interactions, with techniques used indicated. Strikethrough denotes interactions not detected using the respective technique. (B) RDC network localization without NES (left two panels) and with NES (right two panels) (C-F) Pairwise interaction screen of the RDC network using ReLo (left two panels) or nxReLo (right two panels). The conclusions of the tests are indicated next to the images: No, no interaction concluded, yes, interaction concluded, n.c., no conclusion. Question marks denote uncertainty in drawing a conclusion. “Control” refers to the vector shown at the top of the panel, which lacks an insert. The scale bar is 10 µm.

In the absence of an NES, unanchored Cutoff was detectable in both the nucleus and cytoplasm of S2R+ cells, whereas Rhino, Deadlock, Bootlegger, and Moonshiner were predominantly or exclusively nuclear (**Figure 2B**). PH-Cutoff, PH-Rhino, and PH-Moonshiner localized efficiently to the plasma membrane, whereas only a fraction of PH-Deadlock and PH-Bootlegger were membrane-anchored, with the remainder retained in the nucleus (**Supplementary Figure S1**). Upon introduction of the NES, all unanchored RDC network members localized to the cytoplasm (**Figure 2B**), and the membrane anchoring of PH-Bootlegger-NES improved remarkably (**Supplementary Figure S1**). Although PH-Deadlock-NES was now fully cytoplasmic, it still showed suboptimal anchoring (**Supplementary Figure S1**). We resolved this issue by employing mitochondrial surface localization as an alternative to plasma membrane anchoring (see below), a strategy we had previously used (22). Thus, with one exception, the introduction of an NES made all RDC network components (anchored and unanchored) well-suitable for interaction testing in S2R+ cells.

We first screened pairwise interactions systematically among the RDC network, comparing the original ReLo assay with the nxReLo setup. Consistent with the limitations of the ReLo assay, nearly all interaction tests with PH-Bootlegger were inconclusive (**Figure 2C**). In contrast, the nxReLo setup revealed that Bootlegger interacts with Deadlock, but not with Cutoff, Moonshiner, or Rhino. For PH-Cutoff, colocalization with Deadlock in both the nucleus and at the plasma membrane already suggested an interaction in ReLo assays, but with nxReLo this interaction became unambiguous (**Figure 2D**). nxReLo assays further demonstrated that Cutoff does not interact with Bootlegger, Moonshiner, and Rhino, relationships that could not be resolved with the original ReLo assays. Moonshiner did not interact with any other component of the RDC network. While ReLo assays already showed the lack of the Moonshiner - Cutoff interaction, nxReLo confirmed the absence of interactions with the remaining partners (**Figure 2E**). The ReLo assay also showed that PH-Rhino binds Deadlock but not Cutoff (**Figure 2F**), and nxReLo extended this finding by demonstrating that Rhino does not associate with Bootlegger or Moonshiner. PH-Deadlock showed clear interaction with Rhino in ReLo assays and suggested two additional interactions, with Bootlegger and Cutoff, based on strong subcellular colocalization and co-accumulation in nuclear granules (**Figure 3A**). The test with Moonshiner, however, remained inconclusive. Despite suboptimal anchoring of PH-Deadlock-NES in the nxReLo assay, all tests were conclusive, showing that Deadlock interacts with Bootlegger, Cutoff, and Rhino, but not with Moonshiner. Replacing the PH anchor with a mitochondrial localization signal (MLS; see **Methods**) improved the clarity of these results (**Figure 3B**). While MLS-based anchoring did not require an NES, unanchored proteins without an NES did not yield conclusive results (**Supplementary Figure S2**). Despite the improved clarity with an MLS, we prefer plasma membrane anchoring for nuclear proteins, as it is less disruptive to cell morphology.

**Figure 3.**
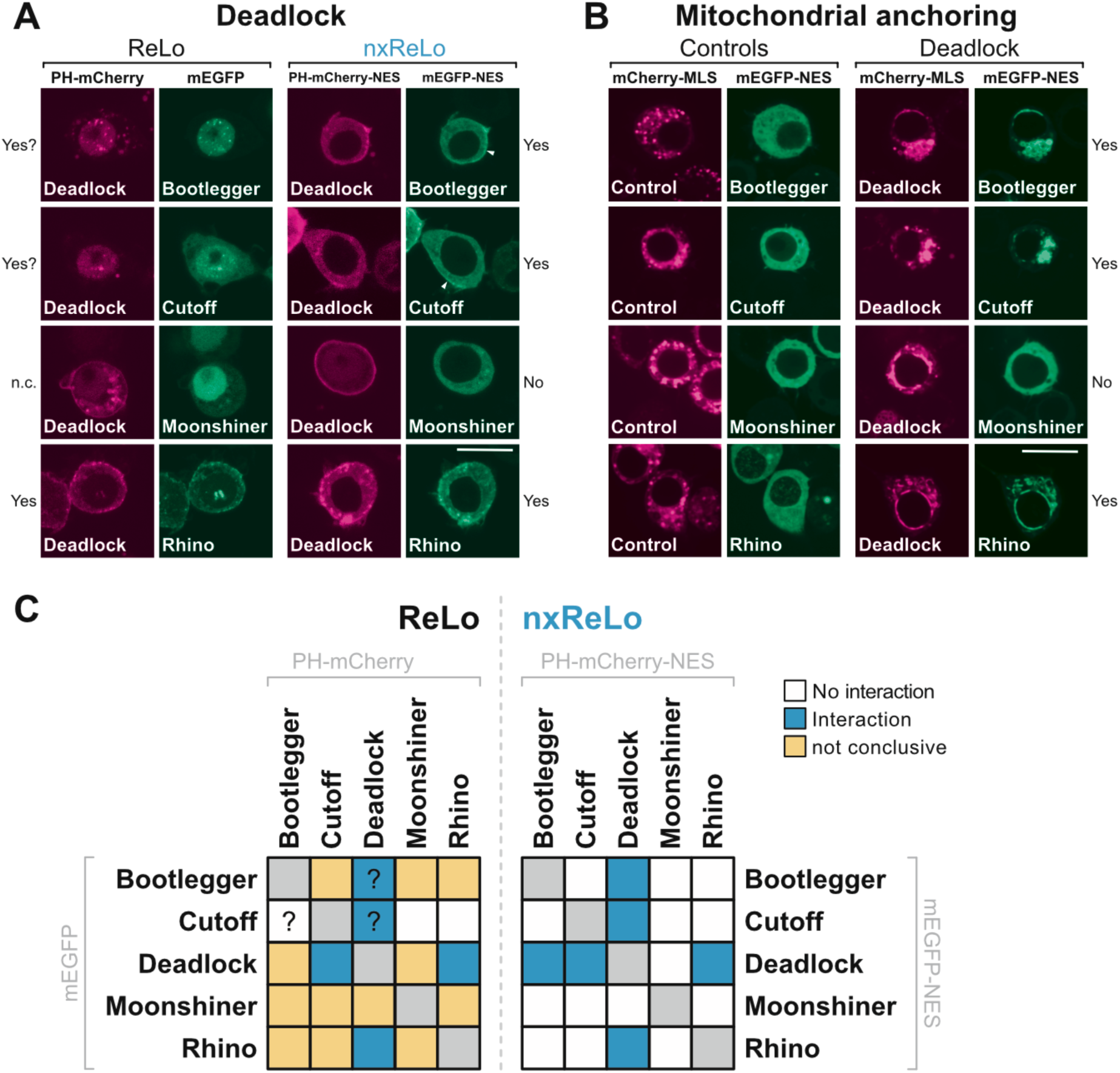
Deadlock interaction screen. (A) Pairwise interaction tests between Deadlock and the rest of the RDC network using ReLo (left two panels) or nxReLo (right two panels). (B) Deadlock was anchored to the mitochondrial membrane via a mitochondrial localization signal (MLS) and tested for pairwise interactions with unanchored Bootlegger, Cutoff, Moonshiner, and Rhino. The conclusions of the tests are indicated next to the images: No, no interaction concluded, yes, interaction concluded, n.c., no conclusion. Question marks denote uncertainty in drawing a conclusion. “Control” refers to the vector shown at the top of the panel, which lacks an insert. The scale bar is 10 µm. (C) Summary of the pairwise interaction screens using ReLo (left) or nxReLo (right). Question marks denote uncertainty in drawing a conclusion.

In summary, the pairwise nxReLo interaction screening between five members of the RDC network enabled conclusive interpretation of all tested interactions (**Figure 3C**). Importantly, the results were independent of the orientation of the protein tagging. nxReLo recapitulated previous findings that Deadlock interacts with Rhino, Cutoff, and Bootlegger (22, 24, 33, 36).

### Characterization of the Cutoff-Deadlock complex

Next, we examined the structurally uncharacterized interactions between Cutoff and Deadlock. Cutoff is related to the Rai1/Dom3Z enzyme family, known to promote transcription termination (37–39). However, Cutoff is a catalytically inactive Rai1/Dom3Z paralog that inhibits co-transcriptional splicing, polyadenylation, and termination of RNA polymerase II transcription at piRNA clusters, ensuring the expression of piRNA transcripts (24, 26, 27). Thus, Cutoff is essential for the expression of piRNA clusters in the heterochromatic regions of *Drosophila* germline cells (32). Similarly, Deadlock is involved in piRNA cluster expression by recruiting the effector protein Moonshiner that activates transcription initiation through TRF2 (22, 23).

Cutoff is composed of a single Rai1-like domain, whereas Deadlock consists of an N-terminal domain (NTD), a long disordered region, and a C-terminal domain (CTD) (**Figure 4A**). Y2H assays have indicated that Deadlock interacts with Cutoff via its C-terminus (aa 661-981) (24, 33). As no structural data is available for the Deadlock-Cutoff complex, we used AlphaFold3 (40, 41) to predict its structure, resulting in a high-confidence model, in which the CTD (aa 695-981) of Deadlock binds to Cutoff (**Figure 4B; Supplementary Figure 3A**).

**Figure 4.**
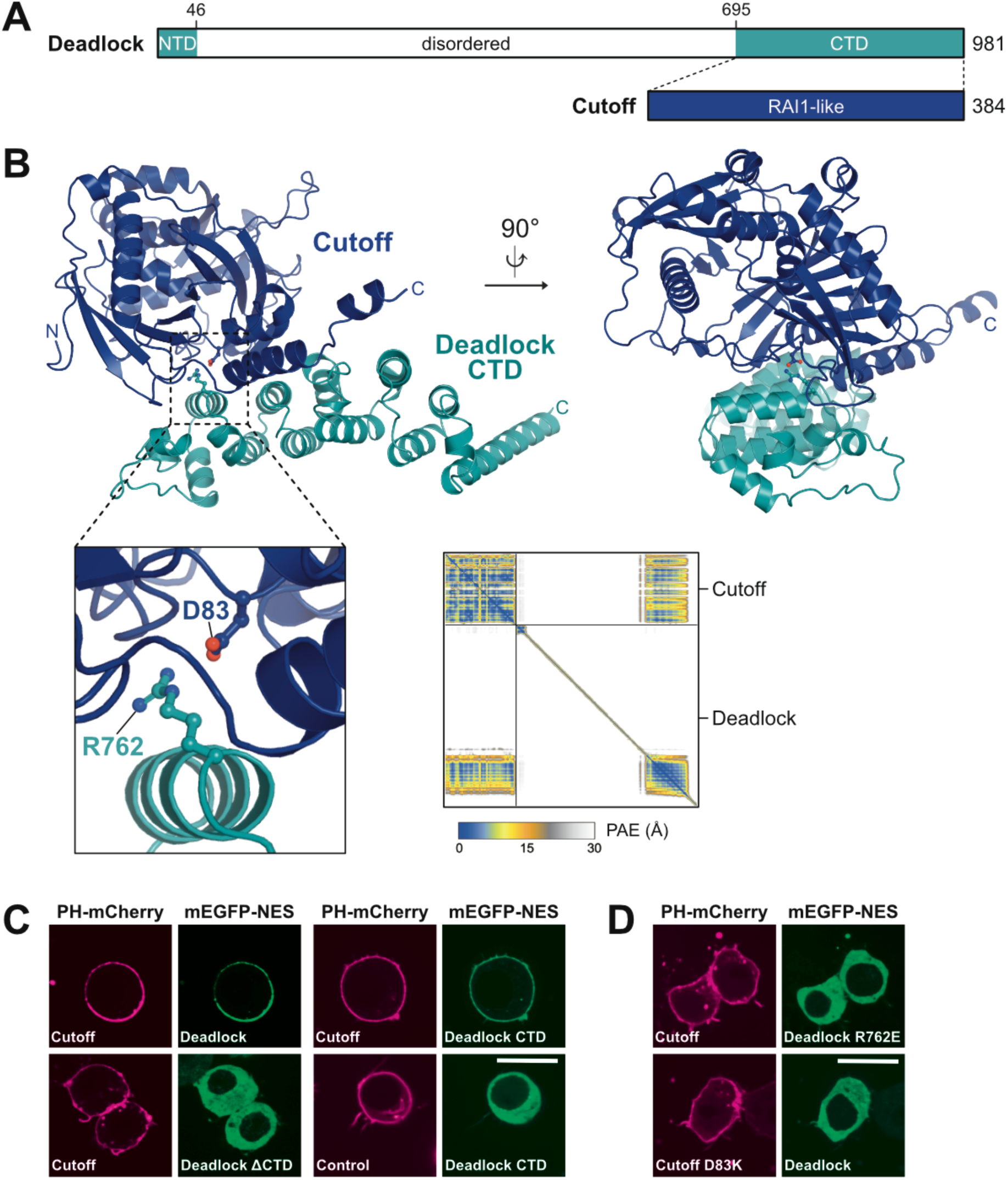
Characterization of the Cutoff-Deadlock interaction. (A) Domain organization of Deadlock and Cutoff. Dotted lines connect the domains involved in the interaction. (B) AlphaFold 3 structure prediction showing Deadlock CTD (aa 700-933; deepteal) binding to Cutoff (dark blue), including a close-up view of the interaction interface, with surface residues tested in mutational analysis indicated in ball-and-stick representation. The Predicted Aligned Error (PAE) plot indicates the high confidence of the predicted model. See also **Supplementary Figure S3A**. (C) Deadlock CTD (aa 695-981) interacted with Cutoff in nxReLo, whereas Deadlock ΔCTD did not. (D) Deadlock R762E and Cutoff D83K point mutations interfered with the Deadlock-Cutoff interaction. “Control” refers to the vector shown at the top of the panel, which lacks an insert. The scale bar is 10 µm.

We validated the structural model using nxReLo. Since Cutoff localized to both the nucleus and the cytoplasm of S2R+ cells, we used Cutoff constructs without an NES in all following experiments. We found that Deadlock lacking the CTD (ΔCTD) did not bind to Cutoff, and that the Deadlock-CTD was sufficient for the interaction with Cutoff (**Figure 4C**). Based on the AlphaFold model, we designed surface mutations at the Deadlock-Cutoff interaction interface (**Figure 4B**). Indeed, neither did the Deadlock point mutant (R762E) bind to Cutoff, nor did the Cutoff mutant (D83K) interact with Deadlock (**Figure 4D**), which is consistent with the model. These results illustrate the easy application of nxReLo to test the binary interaction of a nuclear complex, as well as to map the interaction domains and to identify the interfering point mutations at the interaction interface.

### Characterization of the Bootlegger-Deadlock complex

Using nxReLo and AlphaFold 3, we also investigated the Deadlock-Bootlegger complex (**Figure 5A**), which has not been structurally described either. Bootlegger is recruited to the RDC complex and piRNA clusters through its interaction with Deadlock (33). Bootlegger is involved in the nuclear export of unprocessed piRNA precursors by recruiting the Nxf3-Nxt1 heterodimer and the UAP56 helicase to the piRNA clusters, leading to the export of piRNA precursors by the exportin Crm1 (33, 34).

**Figure 5.**
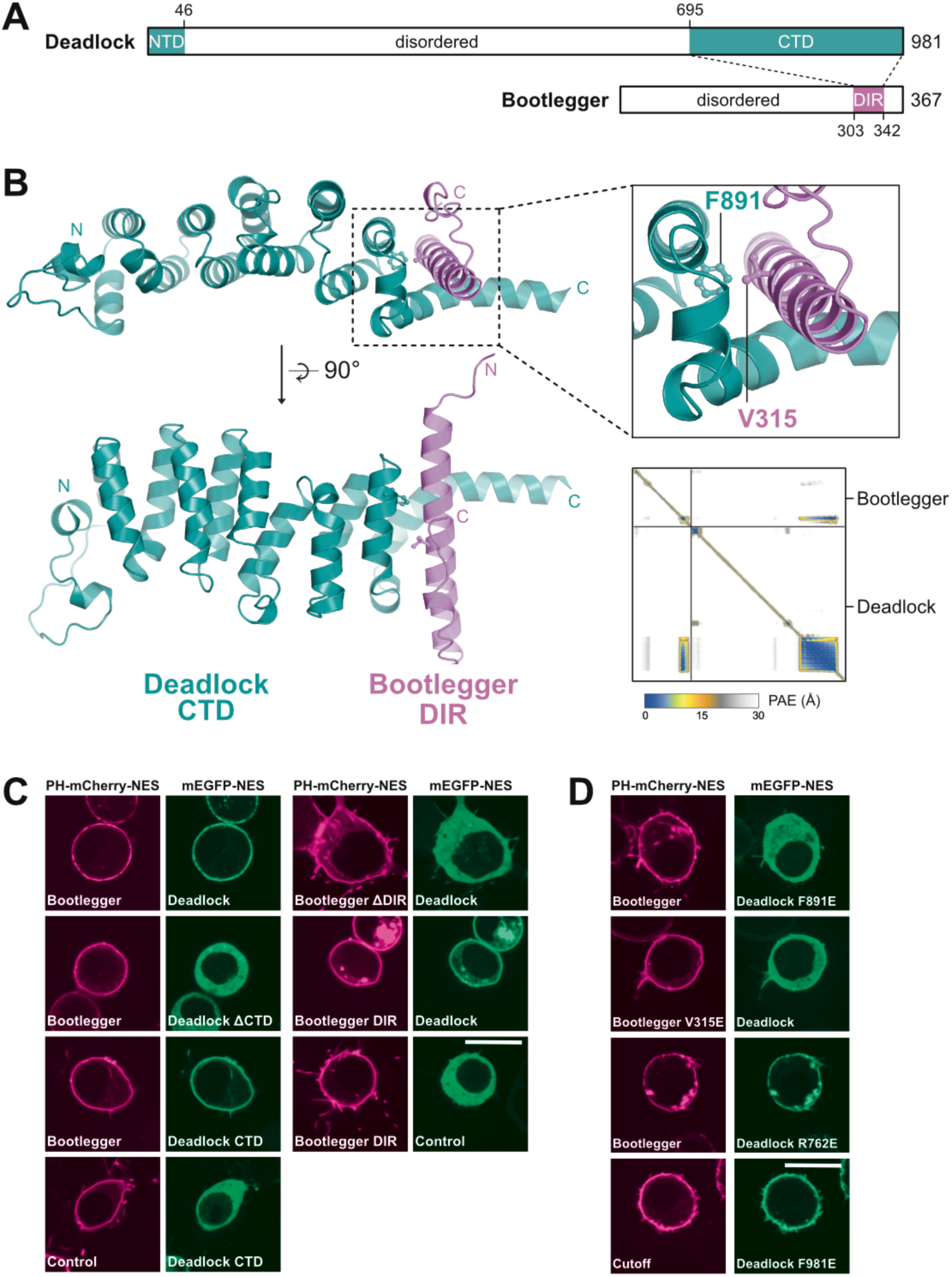
Characterization of the Bootlegger-Deadlock interaction. (A) Domain organization of Deadlock and Bootlegger proteins. DIR, Deadlock interacting region. Dotted lines connect the domains involved in the interaction. (B) AlphaFold 3 structure prediction showing Deadlock CTD (aa 700-933; deepteal) binding to Bootlegger DIR (aa 300-342; pink), including a close-up view of the interaction interface, with residues mutated shown in ball-and-stick representation. The Predicted Aligned Error (PAE) plot indicates the confidence of the predicted model. See also **Supplementary Figure S3B**. (C) In the nxReLo assay, the Deadlock CTD bound to Bootlegger, and the Bootlegger DIR bound to Deadlock. (D) Deadlock F891E and Bootlegger V315E point mutations disrupted the Deadlock-Bootlegger interaction. The Deadlock R762E mutant did not affect the interaction with Bootlegger, nor did the Deadlock F891E mutation affect Cutoff binding. “Control” refers to the vector shown at the top of the panel, which lacks an insert. The scale bar is 10 µm.

Previous Y2H assays have indicated that Deadlock and Bootlegger interact via their C-termini (33). These data are consistent with an AlphaFold3-generated structural model, in which the CTD of Deadlock shows a medium-confidence interaction with the C-terminal α-helix (aa 303-342) of Bootlegger, which we refer to as Deadlock interacting region (DIR) **(Figure 5B; Supplementary Figure 3B)**. Except for this α-helix, Bootlegger is mostly a disordered protein. We confirmed the predicted Deadlock-Bootlegger model using nxReLo assays, which revealed that both the CTD of Deadlock and the DIR of Bootlegger were necessary and sufficient for the interaction (**Figure 5C**). As shown above (**Figure 4**), Cutoff also bound to the CTD of Deadlock, however, at a surface that is distinct from the one bound by Bootlegger.

To interfere with the Deadlock-Bootlegger complex formation, we designed respective mutations in Deadlock (F891E) and Bootlegger (V315E) both of which disrupted the interaction individually (**Figure 5D**). Notably, the Deadlock F891E mutation that interfered with the Bootlegger interaction did not interfere with Cutoff binding, and the Deadlock R762E mutation that disrupted complex formation with Cutoff did not affect the Bootlegger interaction (**Figure 5D**). This demonstrates that both mutations do not affect proper folding of the CTD, and suggests that the CTD of Deadlock has two specific binding sites for the interaction with Cutoff and Bootlegger, respectively.

### Topology of the RDC network

Rhino, Cutoff, and Bootlegger did not directly interact with each other, but each interacted directly with Deadlock (**Figure 6A**), which suggests that Deadlock is an adaptor protein in this complex. To test the capability of Deadlock to bridge interactions within the RDC network, S2R+ cells were cotransfected with three plasmids: two plasmids expressing PH-mCherry and mEGFP fusion proteins, and a third plasmid expressing non-tagged Deadlock. Indeed, we observed an interaction between Rhino, Cutoff, or Bootlegger with any of the other proteins exclusively in the presence of Deadlock (**Figure 6B**). As we did not detect any interactions with Moonshiner in our previous assays (**Figure 3C**), this protein was not included in these bridging assays.

**Figure 6.**
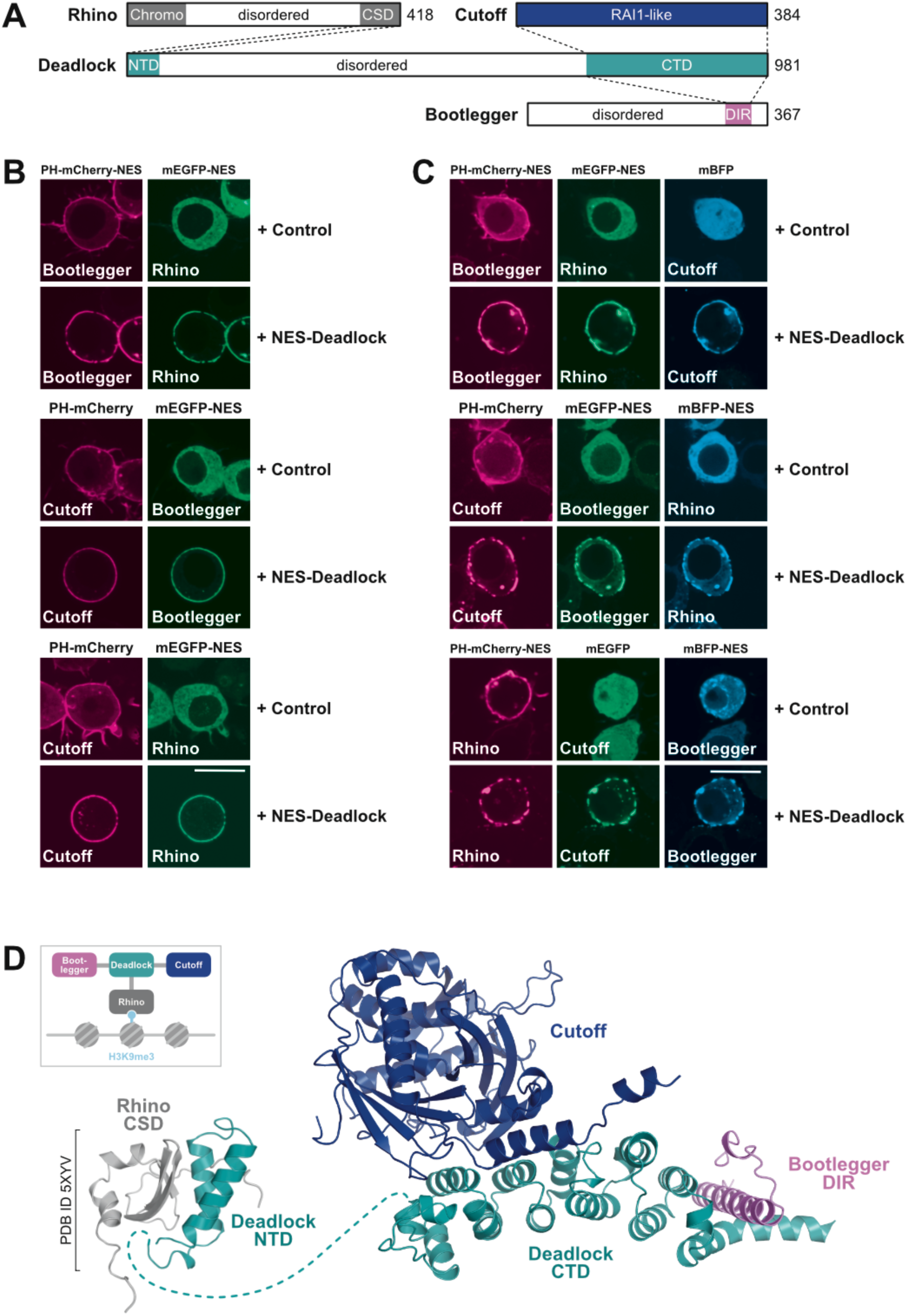
Characterization of the RDC network. (A) Schemes illustrating the mapped interactions within the RDC network. CS, chromo domain; CSD, chromoshadow domain; DIR, Deadlock interacting region. Dotted lines connect the domains involved in the interaction. (B) In nxReLo bridging experiments, PH-mCherry and mEGFP fusions to the proteins indicated were coexpressed with either non-tagged NES-Deadlock or a control vector (pAc5.1A-NES) lacking an insert. After 48 hours, protein localization was analyzed by confocal microscopy. Deadlock could bridge the interaction between the non-interacting proteins of the RDC network. “Control” refers to a vector, which lacks an insert. (C) To bridge and assemble all interacting members of the RDC network via nxReLo, PH-mCherry, mEGFP, and BFP fusions to the proteins indicated were coexpressed with either non-tagged NES-Deadlock or a control vector (pAc5.1A-NES) lacking an insert. After 48 hours, protein localization was analyzed by confocal microscopy. Assembly of the four-component RDC network at the plasma membrane occurred only in the presence of Deadlock. The scale bar is 10 µm. (D) A composite structural model of the RDC network. Deadlock bridges the complex via interactions through its NTD and CTD. Rhino CSD (grey) binds to Deadlock NTD (deepteal), while Cutoff (dark blue) and Bootlegger DIR (pink) bind to Deadlock CTD (deepteal). The dashed line represents the disordered region of Deadlock. Rhino CSD-Deadlock interaction part is taken from (36), PDB: 5XYV. The rest of the composite model was predicted by AlphaFold3.

The previously reported interaction between Cutoff and Rhino based on co-IP (28, 32) was not detected by Y2H (22, 24) or in our pairwise interaction screens (**Figures 2D and 2F**), but was observed in nxReLo assays only in the presence of Deadlock (**Figure 6B**). This bridging could reflect either a simple physical link, with Deadlock binding Rhino via its NTD (36) and Cutoff via its CTD (24) (**Figure 4**), or a conformational change in Cutoff or Rhino induced by Deadlock that enables direct interaction. To distinguish between these possibilities, we tested whether individual Deadlock domains were sufficient to mediate the interaction, but they were not (**Supplementary Figure S4A**), suggesting that the Cutoff-Rhino connection is indirect and requires full-length Deadlock. Consistently, AlphaFold3 did not predict a high-confidence model for the Rhino - Cutoff interaction (**Supplementary Figure S4B**).

Similarly, we did not detect an interaction between Deadlock and Moonshiner (**Figures 2E, 3A and 3B**), despite previous co-IP evidence (23). To explore whether Deadlock’s confirmed partners might induce a conformation that enables Moonshiner binding, we performed nxReLo bridging experiments. However, Moonshiner failed to interact with either the Cutoff-Rhino-Deadlock or the Bootlegger-Rhino-Deadlock complexes (**Supplementary Figure S4C**), again suggesting that the interaction is likely indirect. In line with this, AlphaFold3 did not predict a high-confidence Moonshiner - Deadlock interface (**Supplementary Figure S4D**), and the interaction was not observed in previous Y2H assays (22). However, as we recently observed this interaction between the *Drosophila simulans* orthologs using Y2H (22), the Deadlock - Moonshiner connection might still represent a functionally conserved feature of the RDC network, potentially influenced by context-dependent differences in interaction mode or regulation. Alternatively, since Moonshiner was also found to associate with TFIIA-S and TRF2 in the same co-IP experiments that detected the Moonshiner-Deadlock interaction (23) and in Y2H assays (22), it is possible that these or other unknown factors are required to bridge Moonshiner to Deadlock.

Finally, to further exploit the nxReLo system, we tested interactions between all interacting RDC network members simultaneously by introducing a fourth plasmid expressing a monomeric blue fluorescent protein (mBFP) fusion. In all combinations tested, Deadlock bridged interactions between Rhino, Cutoff, and Bootlegger, assembling the entire complex at the cell membrane (**Figure 6C**). These findings are consistent with earlier results (**Figure 4** and **Figure 5**), indicating that Deadlock binding to Bootlegger and Cutoff is not mutually exclusive but can occur simultaneously. Collectively, these results demonstrate that nxReLo can be used to assemble nuclear multiprotein complexes and study their topology.

## DISCUSSION

Here we developed the nxReLo assay, a fast and simple cell-based method for the investigation of protein-protein interactions designed for its application with nuclear proteins. Using nxReLo, we investigated the interactions between the members of the RDC complex and two associated proteins, a structurally poorly characterized nuclear network required for piRNA transcription from heterochromatin sites (23, 24) and for the export of piRNA precursors (33, 34).

Using nxReLo, we detected pairwise interactions of Rhino, Cutoff, and Bootlegger to Deadlock, but not with other components of the RDC network. Furthermore, a single nxReLo cotransfection experiment showed that all four components can physically assemble into one complex (**Figure 6C**). By combining nxReLo assays with AlphaFold structure prediction and mutational analysis for model validation, we found that Deadlock’s CTD interacts with both Cutoff and the DIR of Bootlegger via two distinct surfaces (**Figure 6D**), providing novel structural insights. Previous studies, including a crystal structure, showed that the NTD of Deadlock binds the C-terminal chromoshadow domain (CSD) of Rhino (36), which recognizes H3K9me3 marks at genomic piRNA loci (24, 28, 35). Together, these findings support a model in which Deadlock links heterochromatic piRNA cluster recognition to Cutoff-mediated transcription antitermination and Bootlegger-dependent export of piRNA precursors (**Figure 6D**). In future studies, the surface point mutations we designed to selectively disrupt the Deadlock-Cutoff and Deadlock-Bootlegger interactions will be valuable tools for dissecting the *in vivo* roles of individual RDC-mediated processes.

The nxReLo assay was designed to test interactions between nuclear proteins within the cytoplasm rather than in their native nuclear environment. While this relocates the proteins from their natural context, if offers two key advantages: first, interactions are less likely to be influenced by endogenous nuclear factors; second, cytoplasmic localization reduces the risk of unintended effects on gene regulation, which could otherwise impact cell viability, especially when working with transcription factors or regulators that are either very potent or not normally expressed in S2R+ cells.

We demonstrated that nxReLo enables the assembly of multi-subunit complexes, a capability not typically addressed by other cell-based interaction assays such as Y2H or bimolecular complementation (BiC) assays. While Y2H can, in principle, accommodate a third protein to test for bridging interactions, negative results are difficult to interpret, as they may simply reflect insufficient expression of the bridging factor. In BiC assays, which rely on the reconstitution of a split fluorescent protein or enzyme (e.g., luciferase or β-galactosidase), interaction detection requires very close spatial proximity of the fusion partners. Although a third component can technically be included, successful bridging may still go undetected if the reconstituted reporter domains are too far apart for complementation, leading to false-negative results.

Similar to other cell-based PPI methods, the use of protein tags in nxReLo may sterically interfere with complex formation. Although we did not encounter this issue in our study, it can often be resolved by altering the tag position. Another challenge in cell-based interaction assays is the testing of proteins that compromise cell viability, such as potent nucleases or proteases. In such cases, toxicity can be mitigated by expressing only functional domains or introducing catalytic site mutations, provided the interaction does not depend on full-length protein integrity or a specific conformation that requires an intact active site. One issue we did encounter with nxReLo is inefficient membrane anchoring of PH-tagged proteins, such as PH-Deadlock. In these cases, we recommend anchoring the bait protein to the mitochondrial surface via an MLS, an alternative approach we successfully applied for Deadlock.

Similar to the original ReLo assay, nxReLo is a qualitative tool designed to efficiently screen and characterize interactions between proteins without assessing their affinity. As previously discussed (18), efficient complex formation, and thus the extent of relocalization, depends on the cellular concentrations of the interacting partners, which are influenced by several uncontrolled factors, including protein expression levels, stability, and subcellular localization. For these reasons, we choose not to quantify the degree of relocalization for a given interaction, as this could lead to the false assumption that it reflects the strength of an interaction.

In conclusion, nxReLo is a simple and effective method for screening and characterizing interactions between nuclear proteins, particularly when recombinant expression for in vitro studies is not feasible or when other cell-based assays fail. Its straightforward setup makes it well-suited for adaptation to high-throughput screening using plasmid libraries, automated microscopy, and image analysis. Although nxReLo is conducted in *Drosophila* S2R+ cells, we have demonstrated the applicability of the ReLo assay to proteins from other species, such as mouse (18), suggesting that also nxReLo may be suitable for studying interactions across diverse organisms. When combined with structure prediction, nxReLo enables rapid structural characterization of nuclear protein complexes. As such, it provides a valuable first step for mapping protein interactions and defining the topology of multi-subunit nuclear complexes. The insight gained can be further complemented by biochemical, structural, and genetic approaches to deepen our understanding of protein complex function and regulation.

## MATERIALS AND METHODS

### Plasmid backbone construction

The plasmids pAc5.1-mEGFP (T6-MJ), pAc5.1-PH-mCherry (HK49) (18), and pAc5.1-mCherry-MLS (JM292) were previously described (22). To generate the pAc5.1-mEGFP-NES (JK298) and pAc5.1-PH-mCherry-NES (JK295) vectors, a modified NS2 NES sequence (TSDEMTKKFGTLTI) (31) was introduced into the pAc5.1-mEGFP and pAc5.1-PH-mCherry vectors C-terminally of the fluorescent tag. For the construction of the non-fluorescent pAc5.1-NES vector (JK301), the pAc5.1 backbone was reverse-PCR-amplified using primers that introduced both a start codon and an optimal Kozak sequence (reverse primer) as well as a linker region (forward primer) upstream of the EcoRV site used for ORF insertion. The modified NS2 sequence was inserted into the amplified vector using an annealed double-stranded DNA oligonucleotide for ligation. We used a modified NS2 NES sequence containing a Val to Ser exchange at the second position, because compared to the wild-type NS2 sequence, it binds less to Crm1 in the absence of RanGTP, facilitating the release of Crm1 from the NES in the cytoplasm (31). We assumed this would reduce the chance of a possible re-import of the NES-containing constructs into the nucleus.

### DNA constructs

Sequences of interest (ORFs) were inserted into the plasmids pAc5.1-mEGFP and pAc5.1-PH-mCherry downstream of the fluorescent tag, and into the plasmids pAc5.1-mEGFP-NES and pAc5.1-PH-mCherry-NES between the fluorescent tag and the NES sequence. In all mEGFP-containing plasmids, the ORFs were inserted into the EcoRV site, and in all mCherry-containing vectors they were inserted into the blunt-end FspAI site. For the construction of the mBFP-containing vectors, the mBFP sequence was amplified from the pCAG-mtagBFP plasmid (42), and the ORFs of interest were inserted via the blunt-end EcoRV site into the pAc5.1 vector using Gibson Assembly (New England Biolabs; E2611L). For anchoring Deadlock to the mitochondria, Deadlock was cloned into pAc5.1-mCherry-MLS upstream of mCherry via the blunt-end FspAI site. For the bridging experiments, Deadlock was inserted into the blunt-end EcoRV site of the pAc5.1-NES vectors. All plasmids containing mutant ORFs were generated by site-directed mutagenesis PCR. Ligation reactions (10 µl) were set up with T4 DNA ligase (Thermo Fisher Scientific), using 50 ng of linearized vector and PCR-amplified insert at a 1:10 - 1:20 molar ratio and containing 5% PEG 4000. Reactions were incubated for 1 h at room temperature. Detailed information on all plasmids used in this study is provided in **Supplementary Table S1**.

### nxReLo assay

*Drosophila* S2R+ cells were cultured at 25°C in Gibco Schneider’s *Drosophila* medium + (L)-glutamine (Thermo Scientific; 21720024) supplemented with 10% fetal bovine serum (Sigma; F9665) and 1 x Gibco Antibiotic-Antimycotic (Thermo Scientific; 15240062). The S2R+ cells were seeded onto a 4-well polymer µ-Slide (Ibidi; 80426) and cotransfected with the desired combination of plasmids using jetOPTIMUS transfection reagent (Polyplus; 101000051). In detail, 600 µl of cells are seeded in each well at a density of 1 × 10^6^ cells/ml, and 61 µl of a transfection mix, including a total of 600 ng of plasmid DNA for cotransfection diluted in 60 µl jetOPTIMUS buffer, and 1 µl of jetOPTIMUS transfection reagent, was added. For cotransfection, 300 ng of each plasmid was used when transfecting two plasmids, 200 ng each for three plasmids, and 150 ng each for four plasmids. The cotransfected cells were incubated at 25°C for 48 h, and images of the live cells were taken with a Nikon Plan Apoλ 100 x NA 1.45 oil objective and a Nikon Ti2-W1 spinning disk confocal fluorescence microscope or a Nikon Apo 60 x NA 1.40 oil-λS objective and a Nikon Eclipse Ti2-AX point scanning confocal fluorescence microscope. The microscopy images were acquired with the NIS Elements AR software and processed using Fiji software (43). The cell(s) shown in the figures are representative of the entire cotransfected cell population.

### AlphaFold structure prediction

All models were generated with AlphaFold 3 (40, 41) using the web interface with standard settings. PAE plots were visualized using UCSF ChimeraX (44) and structures using PyMOL (www.pymol.org).

## Supporting information

Dataset_S1_Cutoff_Deadlock_CTD

Dataset_S2_Bootlegger_DIR_Deadlock_CTD

## ACKNOWLEDGMENTS AND FUNDING SOURCES

We thank Eric Lingren for technical assistance. We are grateful to the constructive feedback on the manuscript from Ralph Grand, Ingrid Lohmann, and Rodrigo Villaseñor Molina. We thank the Nikon Imaging Center at the University of Heidelberg for access to microscopes. We thank the data storage service SDS@hd, supported by the Ministry of Science, Research and the Arts Baden-Württemberg (MWK) and the German Research Foundation (DFG) through the grants INST 35/1314-1 FUGG and INST 35/1503-1 FUGG. This work was funded by the Emmy Noether Program of the German Research Foundation (DFG; JE-827/1-1 and JE-827/1-2 to M.J.), by the Chica and Heinz Schaller Foundation (CHS; 2024 Award to M.J.), by the Novo Nordisk Foundation (NNF18OC0030954 to P.A.) and the Independent Research Fund Denmark (DFF; 9064-00056B to P.A.).

## AUTHOR CONTRIBUTIONS

M.J., P.A. and J.K. designed research; S.I., J.K., and M.M. performed research; S.I., J.K., M.M., P.A., and M.J. analyzed data; and S.I. and M.J. wrote the paper with input from all coauthors.

