## Supplementary Data for "Probing direct interactions between nuclear proteins in cells with nxReLo"

This PDF file includes:

Supplementary Figures S1 to S4
Supplementary Table S1
Supplementary References

Other supporting materials for this manuscript include the following:

*AlphaFold3-generated structural models:*

Dataset S1: Deadlock-CTD - Cutoff complex
Dataset S2: Deadlock-CTD - Bootlegger-DIR complex

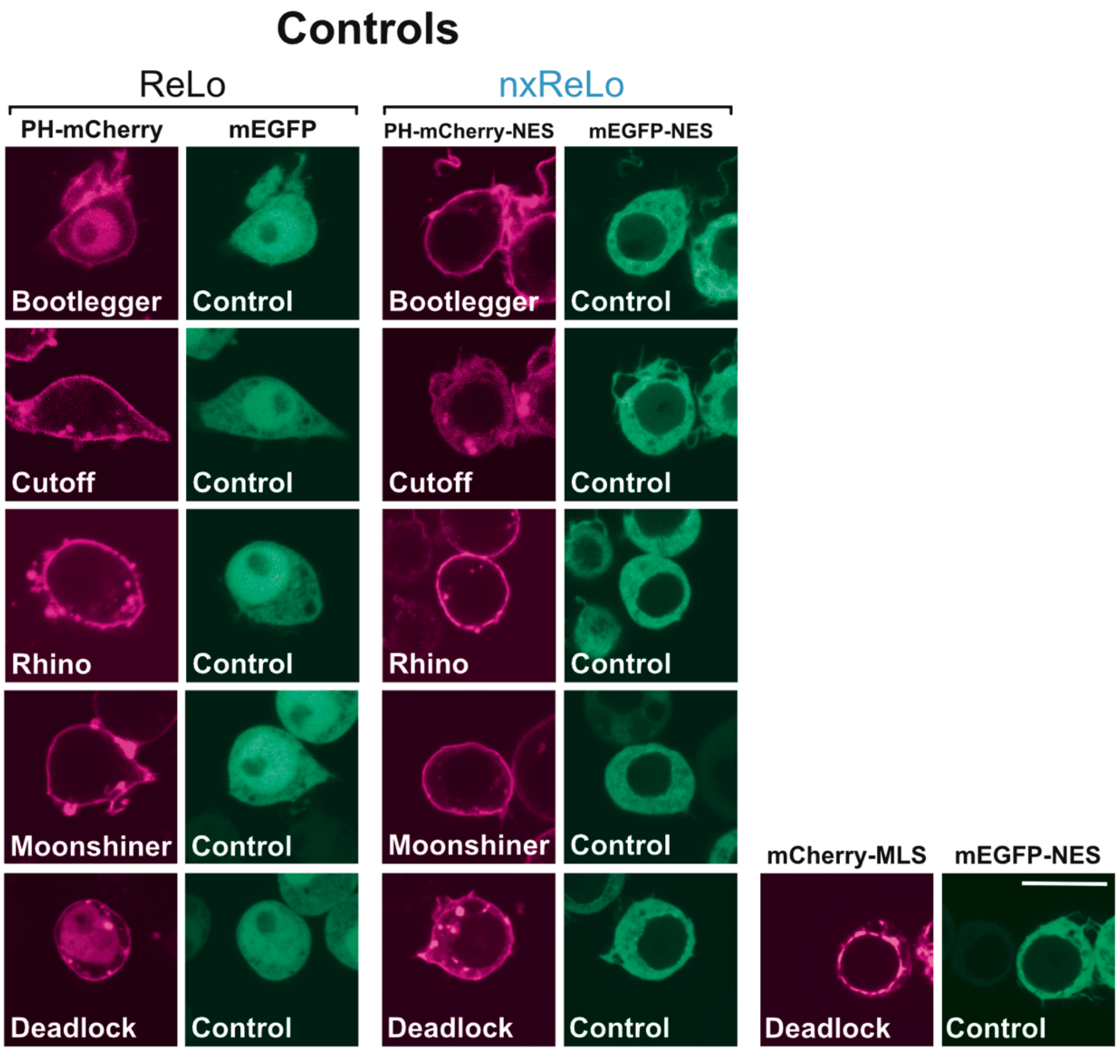


**Supplementary Figure S1. Localization of PH- or MLS-anchored RDC network components.**
Anchoring of the RDC network proteins to the plasma membrane without NES (left two panels) and with NES (right two panels). Alternative anchoring of Deadlock to the mitochondria using an MLS is shown at the bottom right of the figure. “Control” refers to the vector shown at the top of the panel, which lacks an insert. The scale bar is 10 µm.

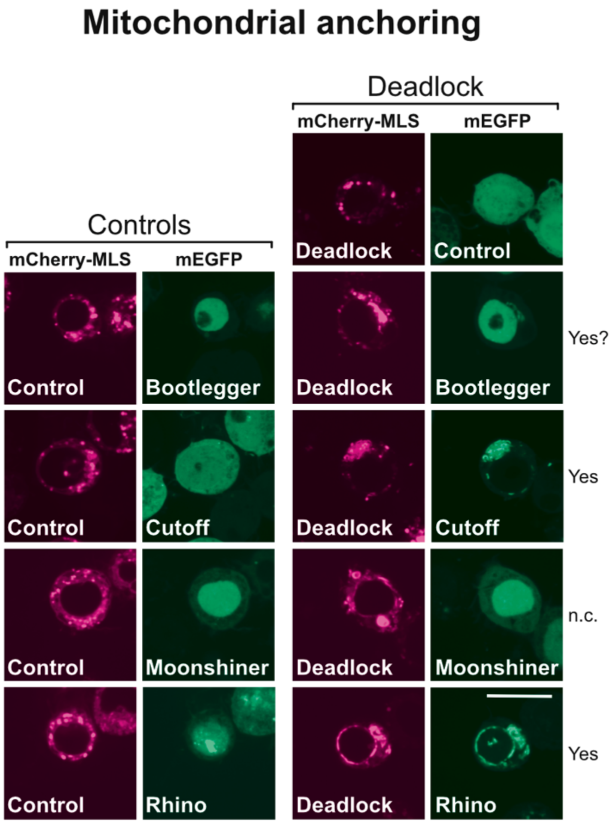


**Supplementary Figure S2. Testing MLS-anchored Deadlock vs. mEGFP-tagged RDC network components lacking an NES.**
The left two panels show the localization of mEGFP-tagged RDC network components in the presence of the control mCherry-MLS vector, which lacks an insert. The right two panels show pairwise interaction tests between mitochondria-anchored Deadlock and other members of the RDC network. The conclusions of the tests are indicated next to the images: No, no interaction concluded, yes, interaction concluded, n.c., no conclusion. The question mark denotes uncertainty in concluding. The scale bar is 10 µm.

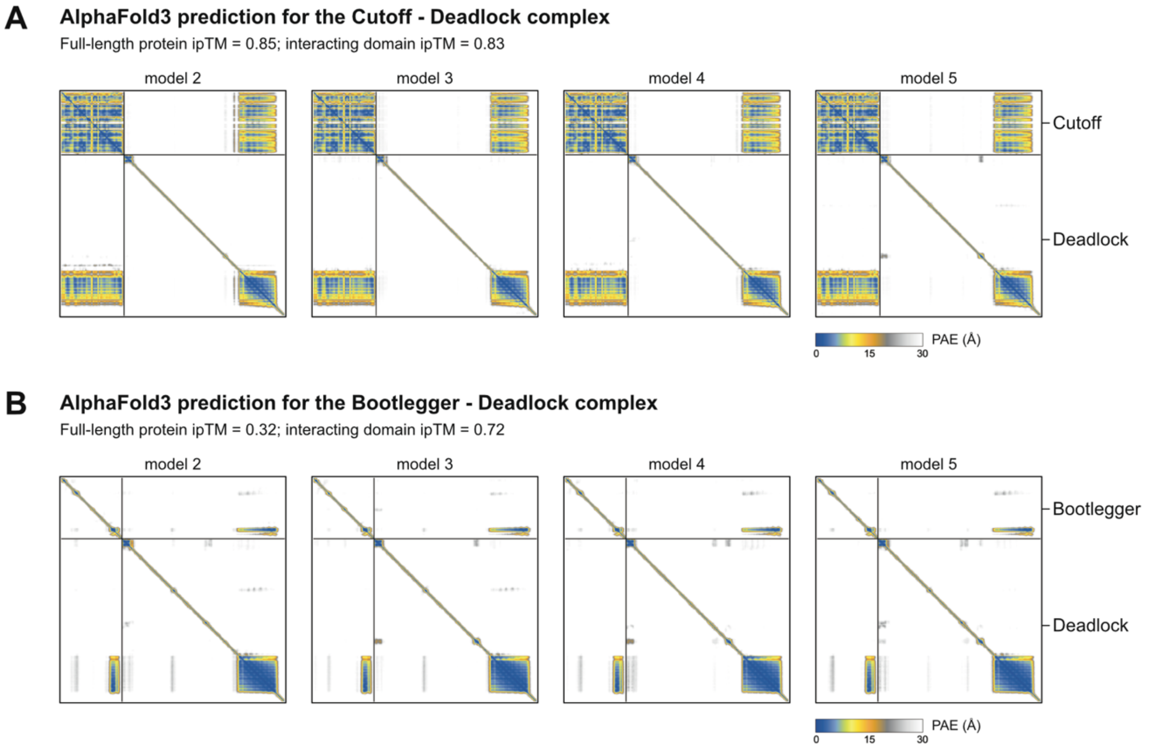


**Supplementary Figure S3. AlphaFold3 predictions.**
The PAE plots for four more AlphaFold3-generated models are shown for the Cutoff - Deadlock (A) and the Bootlegger - Deadlock (B) complexes. The inter-chain predicted template modeling score (ipTM) estimates the confidence in the relative orientation of protein chains within a complex (Evans *et al*, 2021). ipTM values for both the full-length model and for constructs containing only the interacting domains are indicated.

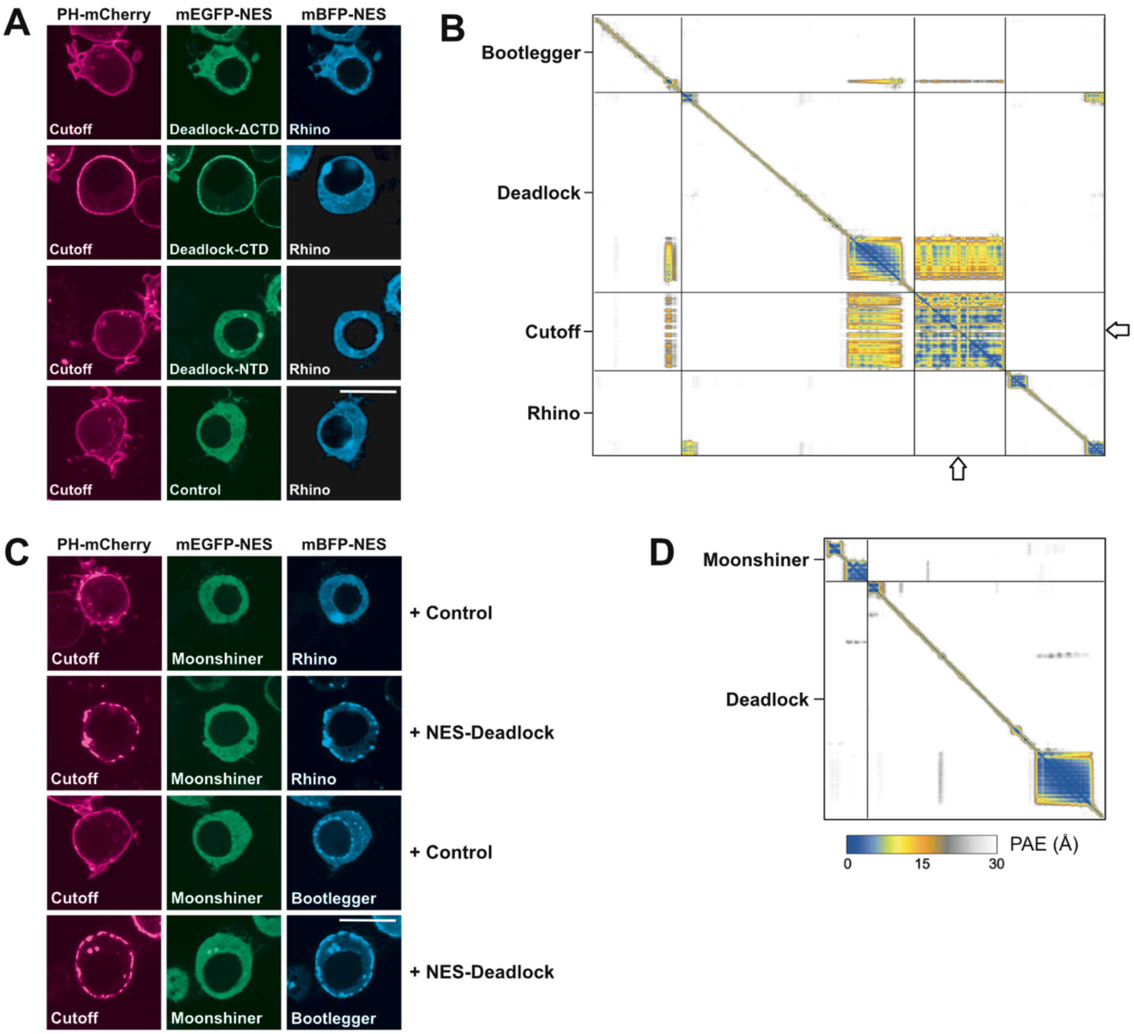


**Supplementary Figure S4. Additional bridging experiments using Rhino and Deadlock.**
(A) In nxReLo bridging experiments, individual Deadlock domains (ΔCTD, CTD, or NTD) fused to mEGFP did not bridge an interaction between Cutoff and Rhino. “Control” refers to the vector shown at the top of the panel, which lacks an insert. The scale bar is 10 µm. (B) PAE plot resulting from an AlphaFold3 structure prediction of a complex composed of Bootlegger, Deadlock, Cutoff, and Rhino showing that no interaction between Cutoff and Rhino is predicted. (C) nxReLo bridging experiments showing that Moonshiner failed to relocalize to the Rhino - Deadlock - Cutoff and Rhino - Deadlock - Bootlegger complexes. “Control” refers to the vector shown at the top of the panel, which lacks an insert. The scale bar is 10 µm. (D) PAE plot resulting from an AlphaFold3 structure prediction showing that no interaction between Moonshiner and Deadlock is predicted.

**Supplementary Table S1.** Plasmids generated and used in this study.
All inserts are *Drosophila melanogaster* sequences. The specific protein isoforms (iso) used are indicated if several exist.

| **Vector** (insertion site) (code) | **Final DNA construct** | **DNA template/ cloning information** | **Code** |
|---|---|---|---|
| **pAc5.1-mEGFP** (EcoRV) (T6-MJ) (Salgania *et al*, 2024) | pAc5.1-mEGFP-**Bootlegger** | *Drosophila* ovarian cDNA | MM33 |
| | pAc5.1-mEGFP-**Cutoff** | *Drosophila* ovarian cDNA | MM35 |
| | pAc5.1-mEGFP-**Deadlock** iso A | *Drosophila* ovarian cDNA | MM3 |
| | pAc5.1-mEGFP-**Moonshiner** | *Drosophila* ovarian cDNA | MM32 |
| | pAc5.1-mEGFP-**Rhino** | *Drosophila* ovarian cDNA | MM38 |
| **pAc5.1-PH-mCherry** (FspAI) (HK49) (Salgania *et al*, 2024) | pAc5.1-PH-mCherry-**Bootlegger** | *Drosophila* ovarian cDNA | EL1 |
| | pAc5.1-PH-mCherry-**Cutoff** | *Drosophila* ovarian cDNA | MM31 |
| | pAc5.1-PH-mCherry-**Cutoff D83K** | Site-directed mutagenesis of MM31 | SI16 |
| | pAc5.1-PH-mCherry-**Deadlock** iso A | *Drosophila* ovarian cDNA | MM37 |
| | pAc5.1-PH-mCherry-**Moonshiner** | *Drosophila* ovarian cDNA | MM29 |
| | pAc5.1-PH-mCherry-**Rhino** | *Drosophila* ovarian cDNA | MM1 |
| **pAc5.1-mEGFP-NES** (EcoRV) (JK298) | pAc5.1-mEGFP-**Bootlegger**-NES | *Drosophila* ovarian cDNA | MM26 |
| | pAc5.1-mEGFP-**Cutoff**-NES | *Drosophila* ovarian cDNA | MM14 |
| | pAc5.1-mEGFP-**Deadlock**-NES iso A | *Drosophila* ovarian cDNA | JK322 |
| | pAc5.1-mEGFP-**Deadlock-CTD**-NES (aa 695-981) | *Drosophila* ovarian cDNA | MM18 |
| | pAc5.1-mEGFP-**Deadlock-ΔCTD**-NES (aa 1-694) | *Drosophila* ovarian cDNA | SI3 |
| | pAc5.1-mEGFP-**Deadlock R762E**-NES | Site directed mutagenesis of JK322 | SI13 |
| | pAc5.1-mEGFP-**Deadlock F891E**-NES | Site directed mutagenesis of JK322 | SI4 |
| | pAc5.1-mEGFP-**Moonshiner**-NES | *Drosophila* ovarian cDNA | MM25 |
| | pAc5.1-mEGFP-**Rhino**-NES | *Drosophila* ovarian cDNA | MM15 |
| **pAc5.1-PH-mCherry-NES** (FspAI) (JK295) | pAc5.1-PH-mCherry-**Bootlegger**-NES | *Drosophila* ovarian cDNA | MM23 |
| | pAc5.1-PH-mCherry-**Bootlegger DIR**-NES (aa 303-342) | *Drosophila* ovarian cDNA | SI7 |

| | | | |
|---|---|---|---|
| | pAc5.1-PH-mCherry-**Bootlegger ΔDIR**-NES (deletion of aa 303-342) | Inverse PCR (deletion) of MM23 | SI6 |
| | pAc5.1-PH-mCherry-**Bootlegger V315E**-NES | Site directed mutagenesis of MM23 | SI8 |
| | pAc5.1-PH-mCherry-**Cutoff**-NES | *Drosophila* ovarian cDNA | MM6 |
| | pAc5.1-PH-mCherry-**Deadlock**-NES | *Drosophila* ovarian cDNA | MM7 |
| | pAc5.1-PH-mCherry-**Moonshiner**-NES | *Drosophila* ovarian cDNA | MM22 |
| | pAc5.1-PH-mCherry-**Rhino**-NES | *Drosophila* ovarian cDNA | JK306 |
| **pAc5.1-NES** (EcoRV) (JK301) | pAc5.1-NES-**Deadlock** iso A | *Drosophila* ovarian cDNA | JK333 |
| **pAc5.1-mBPF-NES** | pAc5.1-mBFP-**Bootlegger**-NES | Gibson assembly | SI18 |
| | pAc5.1-mBFP-**Cutoff** | Gibson assembly | SI19 |
| | pAc5.1-mBFP-**Rhino**-NES | Gibson assembly | SI17 |
| **pAc5.1-mCherry-MLS** (FspAI) (JM292) (Riedelbauch *et al*, 2025) | pAc5.1-**Deadlock**-mCherry-MLS iso A | *Drosophila* ovarian cDNA | JK332 |
